# Human platelet lysate drives clinically compliant generation of vascular mural cells from human pluripotent stem cells

**DOI:** 10.64898/2026.02.03.703640

**Authors:** Laura Yuriko González-Teshima, Wusiman Maihemuti, Kozue Murata, Hidetoshi Masumoto

## Abstract

**Background:** Vascular mural cells (MC) are essential components of vasculature, playing critical roles in tissue regeneration and cell therapy. The use of animal derived ancillary materials, like fetal bovine serum (FBS), in the induction of MC from human pluripotent stem cells (hPSCs), represents one of the biggest limitations to guarantee preclinical safety standards required to use this products in clinical settings. This study aimed to validate human platelet lysate (hPL) as a serum-free alternative for MC differentiation from hPSCs.

**Methods:** Comparison of MC differentiation efficiency from hiPSC using FBS vs hPL supplemented cultures was performed, along with functionality and gene expression assessment through bulk RNA sequencing.

**Results:** Optimization of hPL concentration identified hPL1% as the most effective condition, yielding PDGFR-β+/CNN1+ MC, with a comparable efficiency to FBS10% and similar interaction with endothelial cells in vascular formation assays. However, distinct transcriptional profiles revealed that FBS10% and hPL1% drive differentiation toward different MC subphenotypes; hPL1% promoted contractile gene expression, while FBS10% enriched extracellular matrix pathways. Higher hPL concentrations further shifted differentiation toward cardiomyocytes.

**Conclusion:** In monolayer in vitro differentiation of MC from hiPSC, the differentiation efficiency using hPL 1% supplementation is equivalent to FBS 10%, while supporting a more contractile phenotype. These findings establish hPL as a xeno-minimized, clinically compliant substitute for FBS for hPSC-derived MC differentiation, an important breakthrough for regenerative medicine.

## Background

Reduction of animal-derived products in the differentiation of somatic cell lineages from human pluripotent stem cells (hPSCs) has long been a focus of interest, as the development of xeno-minimized hPSC-derived cell products is an essential safety requirement for their safer application in human cell transplantation therapies[1]. Fetal bovine serum (FBS), a commonly used animal-derived supplement in hPSC differentiation protocols, continues to represent a major limitation in the generation of clinical grade hPSC-derived cell products.

In recent years, human platelet lysate (hPL) has emerged as a promising, serum-free, and clinical grade alternative to FBS across diverse hPSC differentiation protocols[2]. The use of human platelet lysate also offers advantages from the standpoint of medical resource utilization[3–5]. hPL is derived from platelet units that have reached the end of their clinical shelf life and would otherwise be discarded, allowing these materials to be repurposed into a valuable reagent for regenerative medicine[6]. This not only reduces dependency on animal-derived products but also promotes a more sustainable and ethically aligned supply chain. Because hPL is produced from human blood components already processed within established transfusion medicine frameworks[2], its integration into hPSC differentiation protocols provides an additional layer of clinical relevance and regulatory familiarity, further supporting its potential as a translationally suitable supplement [3–5].

Vascular mural cells (MC) are a fundamental cellular component of cardiovascular biology, playing a vital role in the structural and functional architecture of blood vessels, along with endothelial cells[7]. Two subpopulations of MC have been described, mature vascular smooth muscles and pericytes, which are present along different hierarchies of the vascular network[7]. In addition to the essential role of MC in maintaining vascular homeostasis, MCs are known to play a central role in stabilizing tissue architecture when assembling three-dimensional cardiovascular constructs from pluripotent stem cell-derived cell types[8]. Owing to this biological and bioengineering importance, the generation of clinical grade hPSC-derived MC represents a crucial step for advancing cell-based therapies for cardiovascular diseases, with significant implications for the broader field of regenerative medicine[9,10]. Despite the biological and translational importance of MC, progress in establishing clinically applicable differentiation protocols has been uneven. Although great progress has been achieved in the generation of Good Manufacturing Practice (GMP) compliant, serum-free cardiomyocytes[11] and endothelial cells[12], a scalable and clinically compatible animal serum-free differentiation protocol for MC from hPSCs have not yet been reported. Here we aimed to develop a serum-free MC differentiation protocol from human induced pluripotent stem cells (hiPSCs) by replacing FBS with hPL, and evaluated its potential to support efficient and reproducible MC differentiation.

## Methods

### Differentiation of Human iPSC into mural cells

Human iPSCs 201B6 line[13] provided by the Center for iPS Cell Research and Application, Kyoto University was used for all experiments. Maintenance of the hiPSCs was performed as previously described[14,15]. In summary, passage 37 cells were thawed at room temperature with StemFit AK02N medium (AJINOMOTO, Cat No. AK02N, Tokyo, Japan) containing 0.125 μg/cm^2^ iMatrix-511 silk (FUJIFILM Wako Pure Chemical Corp., Cat No. 387-10131, Osaka, Japan) and 10 μM Y-27632 ROCK inhibitor (FUJIFILM Wako, Cat No. 034-24024). After 24 hours, medium change was performed with only StemFit AK02N every other day until 80% confluent. Cells were then dissociated using TrypLE Select (Thermo Fisher Scientific, Cat No. A1285901, Waltham, MA, USA) in 0.5 mM ethylenediaminetetraacetic acid (EDTA)/ phosphate-buffered saline (PBS) solution. Passage 39-41 cells were used for all mural cell differentiation experiments, for which single hiPSCs were seeded at a high density (400,000 cells/cm^2^) onto Matrigel (Corning Life Sciences, Cat No. 356231, NY, USA) pre-coated plates (1:60 dilution) in StemFit AK02N medium with 10 μM Y-27632. Past 24 hours and confirmation of a high density seeding, hiPSCs were covered with a 1:60 Matrigel dilution in AK02N (differentiation day (d) -1). Twenty-four hours later, MC induction protocol was initiated by adding RPMI + B27 medium (RPMI 1640, Thermo Fisher; 2 mM L-glutamine, Thermo Fisher; 1× B27 supplement without insulin, Thermo Fisher) supplemented with 100 ng/mL Activin A (R&D, Cat No. 338-AC-050, Minneapolis, MN, USA) (differentiation day 0). Another 24 hours later (+/-2hours), at d1, the culture was supplemented with 10ng/mL bone morphogenetic protein 4 (BMP4; R&D Cat No. 314-BP-010) and 10ng/mL basic fibroblast growth factor (bFGF; FUJIFILM Wako Cat No. 060-04543) and left for 2 days without further medium changes. At d3, the culture medium was replaced with mural cell induction medium composed of RPMI1640 medium supplemented with 2 mM of L-glutamine and 10% fetal bovine serum (FBS) (Thermo Fisher Scientific, Cat. No. 12483020)[16] or 0,25/0,5/1/3/5/10% human platelet lysate (AventaCell Biomedical Corp, Cat No. HPCHXCRL05, Washington, USA) per tested condition. The culture was then refreshed every other day with the same medium until completing 13 days of culture.

### Tube formation assay

Tube formation assay was performed by seeding a cell suspension (2 × 10^5^ cells/ml) composed of human venous umbilical endothelial cells (HUVEC) (PromoCell, Cat. No C-12200, Heidelberg, Germany) along with either human aortic smooth muscle cells (HAoSMC) (PromoCell, Cat. No C-12533), d13 derived mural cells either cultured with FBS or hPL1% supplementation, were seeded in a 5:1 ratio respectively on a Matrigel pre-coated μ-Slide 15 Well 3D Glass Bottom wells (Ibidi μ-Slide Angiogenesis, Cat. No. 81507. Gräfelfing, Germany) as per manufacturer protocol and incubated at 37ºC, 5%CO2 for 20 hours. A 1:1 ratio of endothelial Cell Growth Medium 2 (EGM-2; PromoCell, Cat. No. C-22011) and smooth muscle cell growth medium 2 (SMGM, PromoCell, Cat.No.C-22062) was used for co-culture of endothelial and mural cell conditions. For HUVEC only EGM-2 was used for culture. All conditions were supplemented with vascular endothelial growth factor (VEGF; FUJIFILM Wako Cat No. 223-01311) 50ng/ml each. Phase contrast images were obtained at 4 and 20 hours of incubation and posteriorly analyzed using Angiogenesis Analyzer plugin for Image J software [17].

### Flow cytometry

Day 13 differentiated mural cells were dissociated with Accutase (Nacalai Tesque, Kyoto, Japan) fir 15 min. Five hundred thousand cells per condition were collected for flow cytometry analysis. Cells were stained with LIVE/DEAD Fixable Aqua LIVE/DEAD Fixable Aqua Dead Cell Staining Kit (Thermo Fisher, Cat No. L34957) in PBS to gate viable cells. Surface staining was performed in flow cytometry buffer (PBS with 5 mM EDTA and 5% FBS) using anti-PDGFRβ conjugated with phycoerythrin (PE) (clone 28D4, 1:100; BioLegend, Cat No. 558821 Franklin Lakes, NJ, USA) for MC, anti–VE-cadherin– PE (clone 55-7H1, 1:100; BioLegend Cat No. 560410) for endothelial cells, and anti-CD90–APC (clone 5E10, 1:100; BioLegend Cat No. 328114) for stromal cells identification. For intracellular staining, cells were fixed with 4% PFA and permeabilized in PBS with 5% FBS and 0.75% saponin (Nacalai tesque, Cat No. 30502-42 Kyoto, Japan), then stained with anti–cardiac troponin T (cTnT, clone 13-11; Thermo Fisher Cat No. MS-295-P0) labeled with allophycocyanin (APC) using Zenon technology (1:50; Thermo Fisher, Cat No. Z25051). Samples were analyzed on a CytoFLEX S flow cytometer (Beckman Coulter, Brea, CA, USA), acquiring 10,000 events per condition, and data were processed with CytExpert software (Beckman Coulter).

### Immunofluorescence Staining

For single-cell day 13 mural cell immunofluorescent staining, cells were separately seeded (5000 cells/well) per condition into IBIDI μ-Slide 15 Well 3D Glass Bottom wells (Ibidi μ-Slide Angiogenesis, Cat. No. 81507, Gräfelfing, Germany) and cultured for 24 hours in fresh day 13 medium and then fixed in 4% PFA for 20 minutes at room temperature. Cells were rinsed with Dulbecco’s phosphate-buffered saline (DPBS) and stored at 4 °C until staining. Immunohistochemical staining was performed using primary antibodies goat anti–alpha-smooth muscle actin (1:200; Abcam, Cat. No. ab21027, Cambridge, UK), rabbit anti–PDGFRβ (1:100; Invitrogen, Cat. No. MA5-24885, Waltham, MA, USA), and mouse anti–calponin 1 (1:100; R&D Systems, Minneapolis, MN, USA), followed by incubation with the respective secondary antibodies: donkey anti-goat IgG (Invitrogen, Cat No. A32849TR), donkey anti-rabbit IgG (Invitrogen, Cat No. A-31572) and donkey anti-mouse IgG (Invitrogen, Cat No. A-21202). Tube formation assay samples were stained with mouse anti–calponin 1 (1:100; Invitrogen, Cat. No. MA5-11620), goat anti–VE-cadherin (1:100; Abcam, ab33168), and rabbit anti–PDGFRβ (1:100; Abcam, ab313777), followed by incubation secondary antibodies donkey anti-goat IgG (Invitrogen, Cat No. A-21432), donkey anti-mouse IgG (Invitrogen, Cat No. A-21202), and donkey anti-rabbit IgG (Invitrogen, Cat No. A-31573). Clearing was performed for tube formation samples with RapiClear® 1.49 (SunJin Lab Co., Cat. No. RC149001, Hsinchu City, Taiwan). All imaging was conducted on a Leica DMi8 widefield microscope (Leica Microsystems, Wetzlar, Germany) and processed using Imaris, version 10.1 (Bitplane, an Oxford Instruments Company, Zurich, Switzerland). Each result was verified by at least two independent experimental repeats.

### Bulk RNA sequencing

Total RNA was extracted from differentiation day 13 cultured cells using RNeasy Mini Kit (Qiagen, Cat No. 74104; Hilden, Germany), quality checked with NanoDrop One Microvolume UV Spectrophotometer (Thermo Fisher Scientific, Waltham, MA, USA) and stored at -80ºC until sequencing. Bulk RNA sequencing was commissioned to CiRA Foundation. In brief, 200 ng total RNA used for library using Illumina TruSeq Stranded mRNA. Sequencing was performed using the NovaSeq Reagent Kit v1.5 with a 101-8-8-101 cycle. The resulting FASTQ data was used for differentially expressed gene analysis. GraphPad Prism 10.6.0 software (GraphPad Software, Inc., Boston, MA, USA) was used for heatmap illustration generation. GO enrichment analysis was performed using SHINY GO for the upregulated and downregulated genes. [18] R package clusterProfiler was used to perform MA plot analysis.

### Statistical analysis

GraphPad Prism 10.6.0 software (GraphPad Software, Inc., Boston, MA, USA) was used for all statistical analysis. All included data variables are continuous and analysis are presented by medians with 95%IC unless stated otherwise. Normality was proved by visual boxplot distribution analysis and Shapiro-Wilk test. Parametric statistical tests were applied. For flow cytometry data two-way ANOVA with multiple comparison analysis correction was applied to compare cell population per differentiation condition. Furthermore, for the tube formation assay one way ANOVA with post-hoc Tukey’s multiple comparison analysis was performed. An alpha value 0,05 was set for all conditions.

## Results

### Optimization of hPL concentration for mural cell differentiation protocol in 2D monolayer culture

Using a previously established mural cell differentiation method[19], mural cells were generated using a 13-day protocol, with either FBS 10% or differential concentrations of hPL supplementation (Figure 1a). MC differentiation efficiency on day 13 showed clear variation according to hPL concentration compared to FBS 10% (Figure 1b). Supplementation of hPL 1% proved to be as efficient as FBS 10% for MC [platelet derived growth factor receptor beta (PDGFR-β) positive cells] differentiation culture, with no significant difference between the differentiation percentage of the two groups (p=0.7906) (Figure 1b). Between hPL concentrations, hPL 1% had the highest MC differentiation efficiency, closely followed by hPL 3% (Figure 2a and 2b). Although no statistically significant difference was observed between hPL1 and 3% groups (p=0.9955) (Figure 2b), hPL 1% exhibited the highest median (50^th^ percentile) value for both cell count and viability in all hPL concentrations (Figure 1c and 1d, respectively).

**Figure 1.**
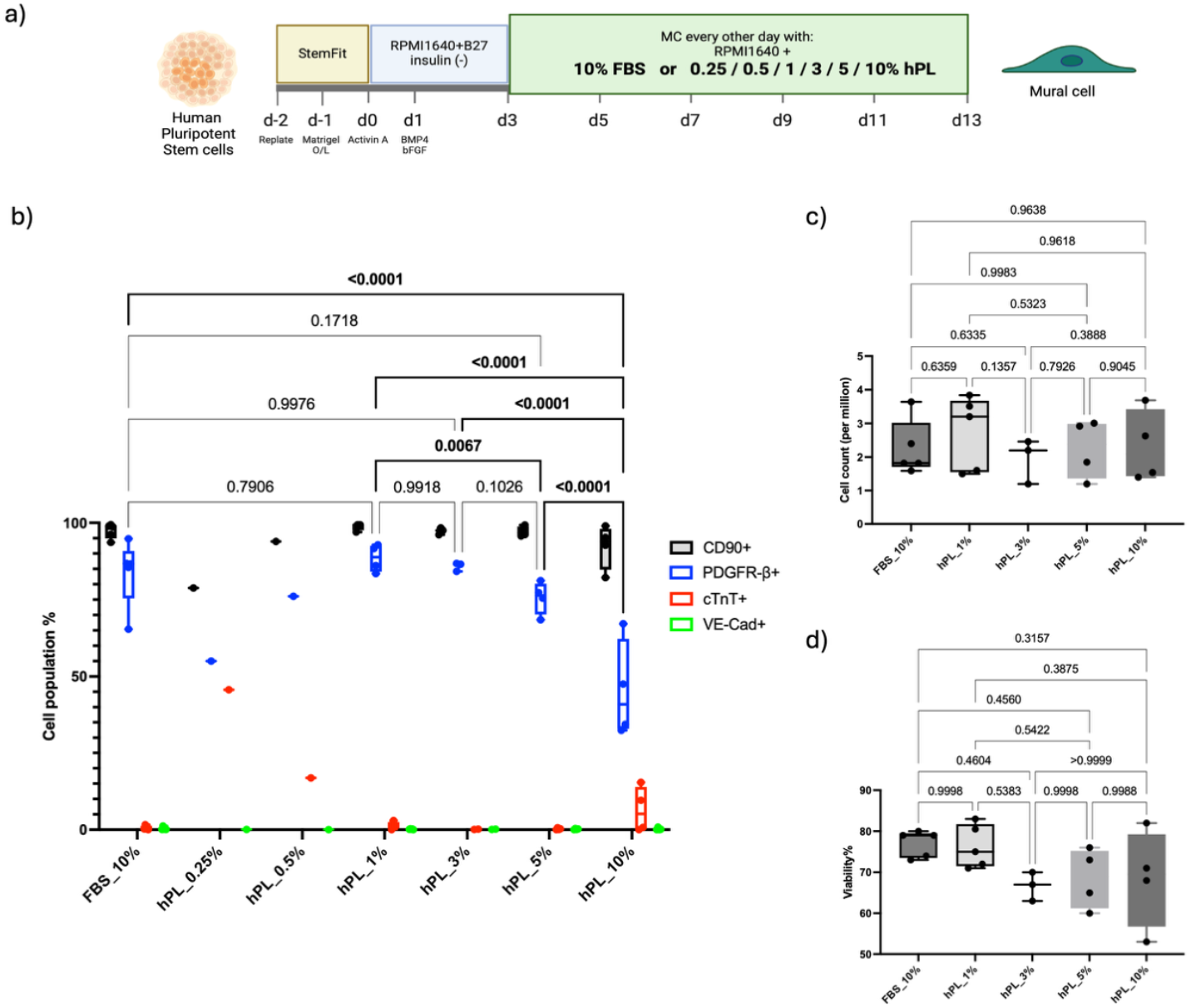
Optimization of human platelet lysate concentration for vascular mural cell differentiation protocol in 2D monolayer culture. **a)** Schematic representation of the 13-day differentiation protocol from 201B6 human pluripotent stem cell (hiPSC) line into vascular mural cells. **b)** Vascular mural cell differentiation efficiency at day 13 per human platelet lysate (hPL) differential concentration and fetal bovine serum (FBS) supplemented conditions (median +/-95% CI). CD90+: Cluster of Differentiation 90 positive cells, representing stromal cell population; PDGFR-β+: platelet derived growth factor receptor beta positive cells, representing vascular mural cell population; cTnT+: Cardiac troponin T positive cells, representing cardiomyocyte population; VE-Cad+: VE-cadherin (CD144) positive cells, representing endothelial cell population. Statistical analysis for PDGFRβ expression was performed using two-way ANOVA with multiple comparison analysis. Significant differences (p<0,05) are indicated by **bold** parenthesis. Data for hPL 0.25% and 0.5% represent n=1 and were excluded from statistical analysis. All other conditions have a minimum of n=3 replicates. **c)** Total cell count (n≥3) and **d)** cell viability at day 13 of differentiation in hPL and FBS supplemented culture conditions (n≥3).

**Figure 2.**
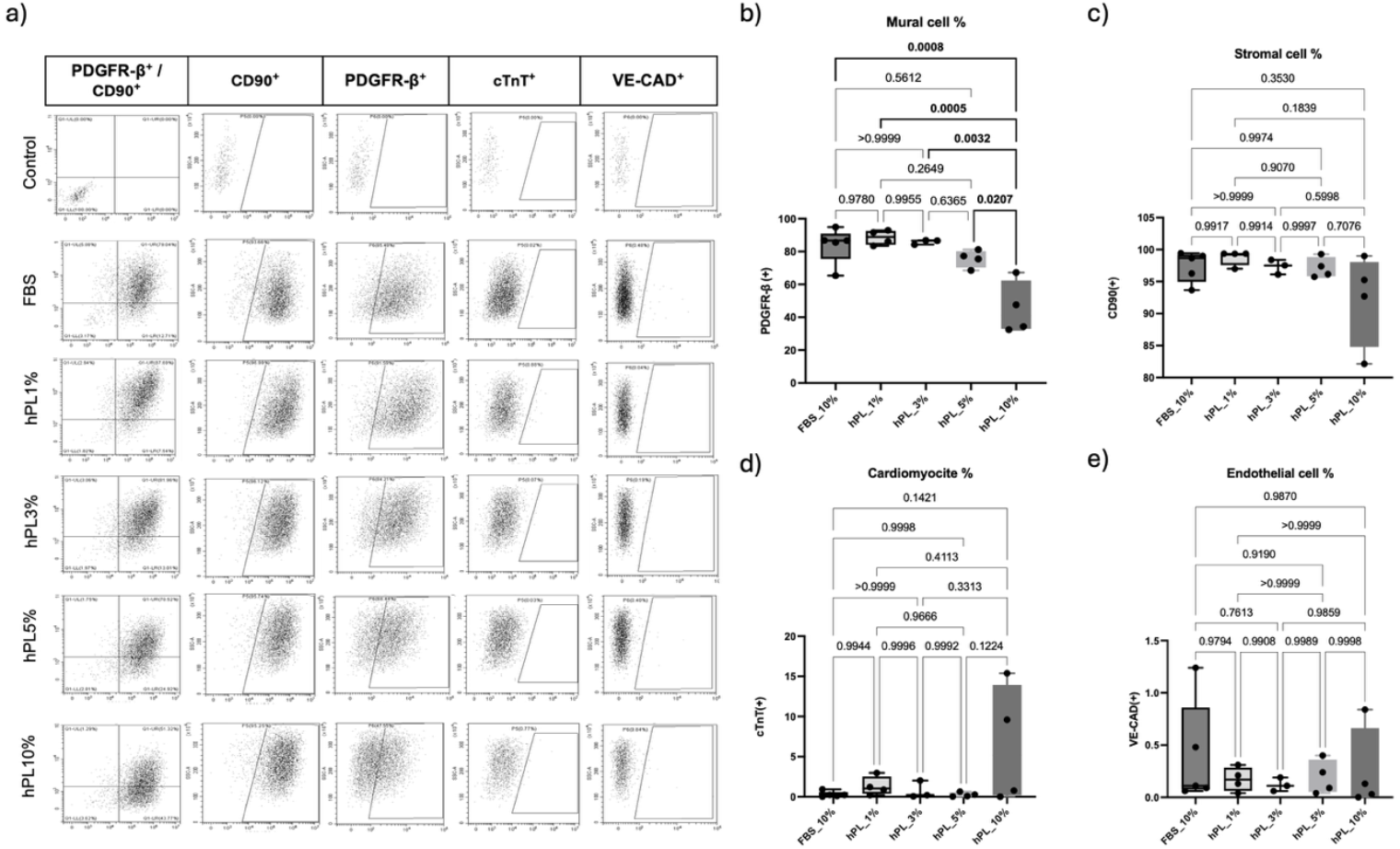
Flow cytometry characterization of human pluripotent stem cell (hiPSC) derived cell populations at day 13 of differentiation. **a)** Representative gating strategy for flow cytometry characterization of hiPSC derived cell populations at day 13 of differentiation. Vascular mural cells are identified as platelet-derived growth factor receptor beta positive (PDGFR-β^+^); Stromal cells as cluster of differentiation 90 positive cells (CD90^+^), cardiomyocytes as cardiac troponin T positive (cTnT^+^) and endothelial cells as VE-cadherin positive (VE-CAD^+^) (n≥3). **b)-e)** Differentiation efficiency of **b)** mural cells, **c)** stromal cells, **d)** cardiomyocytes and **e)** endothelial cells at day 13 under different hPL concentrations and FBS supplemented conditions. Data shown as box and whiskers plots with median, interquartile range and min-max values. Statistical analysis was performed using two-way ANOVA with multiple comparisons (n≥3). Significant differences (p<0,05) are indicated by **bold** parenthesis.

For increasing concentrations above hPL 1%, the differentiation efficiency of MC decreased (Figure 1b, 2a and 2b). Specifically hPL 10% supplementation showed a shift in the differentiated cell population composition at day 13, showing a significant decrease in the mural (Figure 2b) and CD90 positive stromal cell (Figure 2c) population, at the expense of an increase in the percentage of cardiomyocytes up to around 10%, along with a slight increment of VE-cadherin (CD144) positive endothelial cell differentiation (Figure 2d and 2e, respectively), although the variation was quite high between replicates. Furthermore, even though all differentiation conditions looked quite homogeneous along the 13 days of differentiation (Figure 3a), immunofluorescence analysis of MC specific markers showed differentially enhanced fluorescence of pericyte marker PDGFR-β, vascular smooth muscle cell (vSMC) markers anti alpha smooth muscle actin (ACTA2) and calponin 1 (CNN1) for hPL 1% derived MC cells at day 13 compared to all other hPL conditions and even FBS (Figure 3b). Because of the above, hPL 1% supplementation was selected as the optimal concentration to compare with FBS 10% xeno-supplemented standard culture for MC differentiation efficiency.

**Figure 3.**
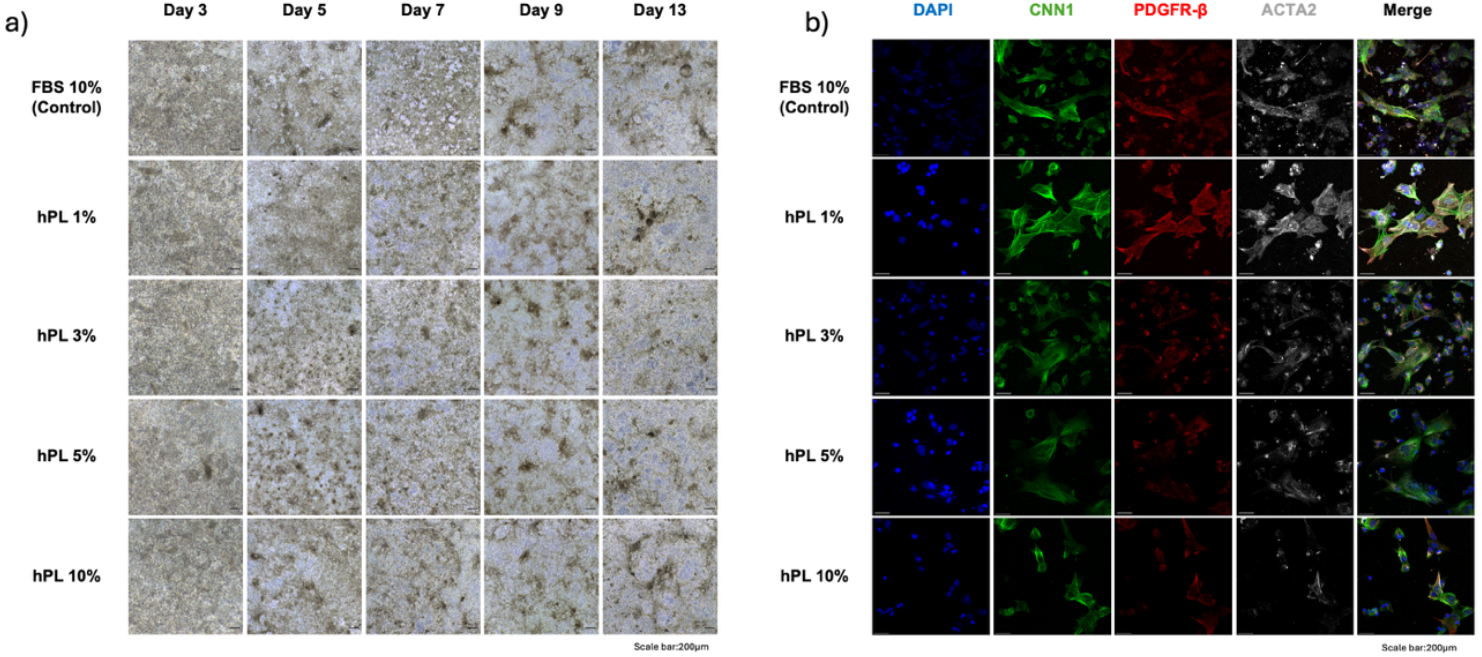
Morphological and immunofluorescent characterization of human pluripotent stem cell (hiPSC) derived vascular mural cells. **a)** Phase contrast representative images showing cell culture morphology across conditions from day 3 to 13 of the mural cell differentiation protocol with no significant differences observed between conditions (n≥3). FBS: fetal bovine serum. hPL: human platelet lysate. **b)** Representative immunofluorescent analysis of day 13 hiPSC derived cell population, showing expression of mural-specific markers under each culture condition (n=3). FBS: fetal bovine serum. hPL: human platelet lysate. DAPI: 4′,6-diamidino-2-phenylindole. CNN1: Calponin 1. PDGFR-β: platelet-derived growth factor receptor beta. ACTA2: α-smooth muscle actin.

### Transcriptomic profiling of hiPSC-derived mural cells generated with hPL

As FBS 10% and hPL 1% supplemented cultures retrieved similarly efficient percentages of MC population with positive markers both in cytometric and immunofluorescence analysis, further evaluation of the genetic profile and functionality of the differentiated mural cells was carried out. Bulk RNA sequencing was performed on both FBS 10% and hPL 1% derived cell populations, as well as hPL 10% due to the increased cardiomyocyte cell differentiation. Principal component analysis showed significant transcriptional differences between both FBS and hPL conditions, and between both hPL 1% and 10% cell populations. FBS and hPL 1% had a shorter Euclidean distance compared to hPL 10% with both above; indicating FBS and hPL 1% have a closer genetic expression compared to hPL 10% as evidenced in both the cytometric and immunofluorescent analysis (Figure 2b, 3b and 4a). Functional analysis of hPL 1% showed significant enhancement of gene ontology (GO) molecular pathways related to structural constituent of muscle, actin and cytoskeletal binding, calcium ion binding, and overall receptor signaling activity compared to FBS (Figure 4b). On the contrary, pathways related to extracellular matrix constituent and regulation, as well as transmembrane transported activity are downregulated in hPL 1% compared to FBS, indicating a clear difference in the molecular functionality profile of the hPL1% vs FBS 10% derived MC (Figure 4c). This was further confirmed by analyzing the differential expression of mural cell-related genes of interest.

**Figure 4.**
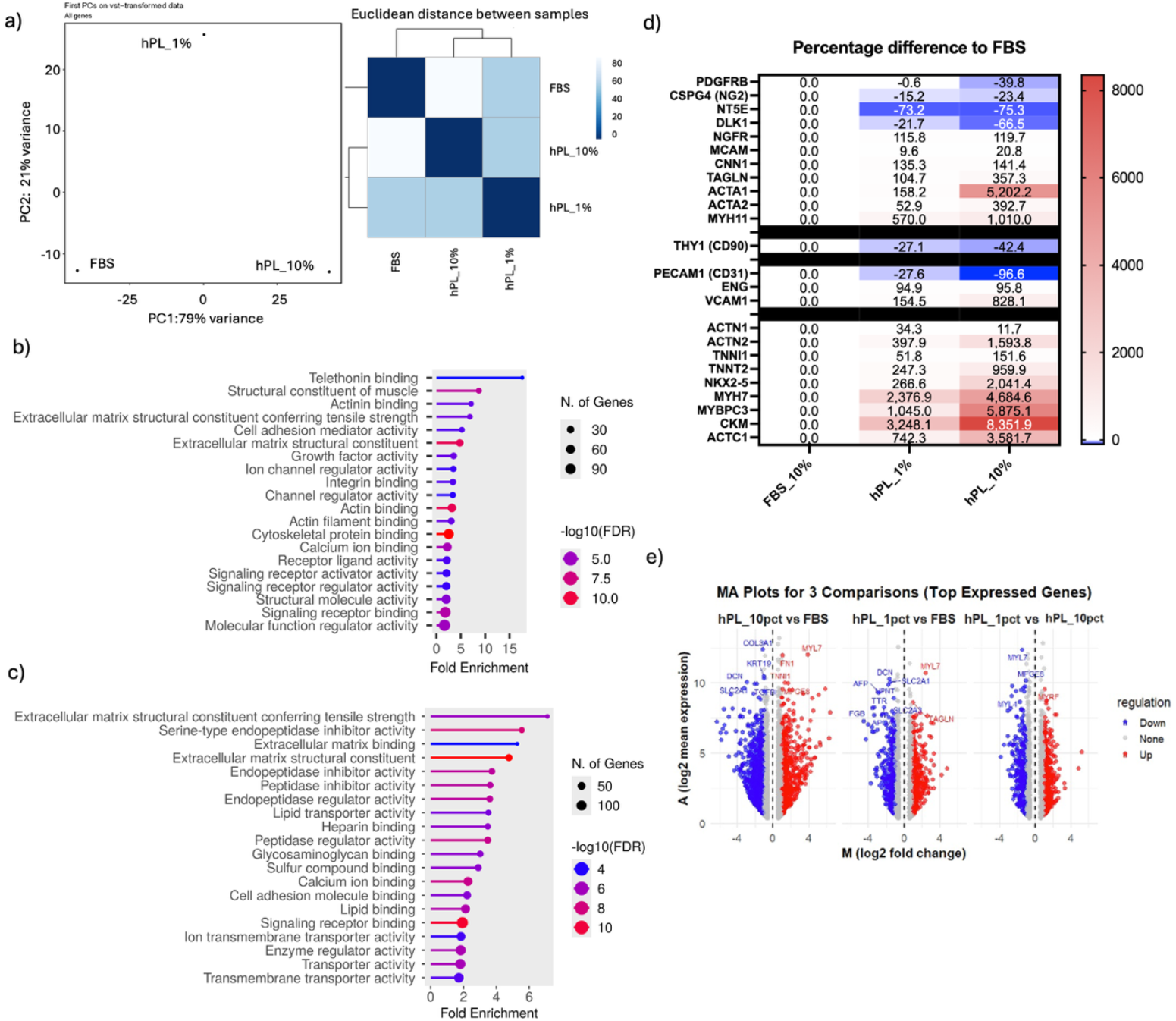
Transcriptomic profiling of hiPSC-derived mural cells generated with hPL evaluated by bulk RNA-sequencing analysis. Principal component analysis (PCA) and Euclidean distance of day 13 hiPSC derived mural cells. Gene ontology enrichment analysis of differentially expressed genes upregulated in hPL1% condition compared to FBS. **c)** Gene ontology enrichment analysis of differentially expressed genes downregulated in hPL1% condition compared to FBS. **d)** Heatmap showing the percentage difference in gene expression relative to FBS-supplemented conditions, associated with mural cells, endothelial cells, and cardiomyocytes (separated into respective blocks in the indicated order). **e)** MA plot of gene expression changes between three different comparison groups hPL10% vs FBS, hPL1% vs FBS, hPL 1% vs hPL10% 1. Red and blue dots represent significantly upregulated and downregulated genes (|log2FC| ≥ 1, FDR < 0.05), respectively. Grey dots indicate non-significant genes.

Although both FBS and hPL 1% seemed to have similar increased expression of PDGFR-β over hPL 10% derived cells, which showed a 39.8% reduced expression compared to FBS (Figure 4d), other recognized pericyte associate markers like NG2, DLK1 and NT5E seemed to be under expressed in both hPL conditions compared to FBS. Nevertheless, the sustained expression of PDGFR-β was concomitantly observed with an interesting increased expression of genes related to smooth muscle contractile machinery like calponin 1 (CNN1), alpha actin smooth muscle (ACTA2), myosin heavy chain 11 (MYH11) and transgelin (TAGLN) [20–22] in hPL1% by more than 100% compared to FBS (Figure 4d). TAGLN and MYH11 were one of the top differentially upregulated genes on hPL1% condition vs FBS (Figure 4e) suggesting a phenotype closer to contractile vSMC in hPL 1% condition derived cells. For hPL 10%, increased ACTA2 and MYH11 expression by more than three and almost ten times respectively, compared to the other conditions, was also associated with significantly increased expression of cardiomyocyte markers like TNNI1, TNNT2, ACTN2, MYBPC3, CKM, ACTC1, MYH7, NKX2-5 (Figure 4d and 4e), which correlates with the increase cTnT positive population observed in the flow cytometry analysis (Figure 2d). Furthermore, hPL1% showed a slightly higher cardiomyocyte related gene expression like MYL7 as a top upregulated gene (Figure 4e), although much less than hPL10% but higher than FBS condition; suggesting at lower concentration hPL might favor both MC and cardiomyocyte induction pathways. Endothelial marker (CD31) was overall under expressed in all conditions and no great differences were observed with stromal cell markers (Figure 4d).

### Functional characterization of hiPSC derived mural cells generated with hPL

Finally, we compared the functionality of the MC differentiated in both cultures supplemented with FBS or hPL1%, by assessing their differential modulation of endothelial cell angiogenic potential though a tube formation assay (Figure 5a). Both FBS and hPL 1% derived MC cocultures with HUVEC showed reduced angiogenic modulation compared to positive control, represented by human aortic smooth muscle cells (HAoSMC), as evidenced by shorter total angiogenic network and branching length (p<0,01) (Figure 5b and 5c respectively) and decreased network complexity (p<0,001) (Figure 5d). These values were equivalent to negative control (HUVEC only), suggesting functional immaturity of MC derived from FBS and 1%hPL cultures compared to HAoSMC.

**Figure 5.**
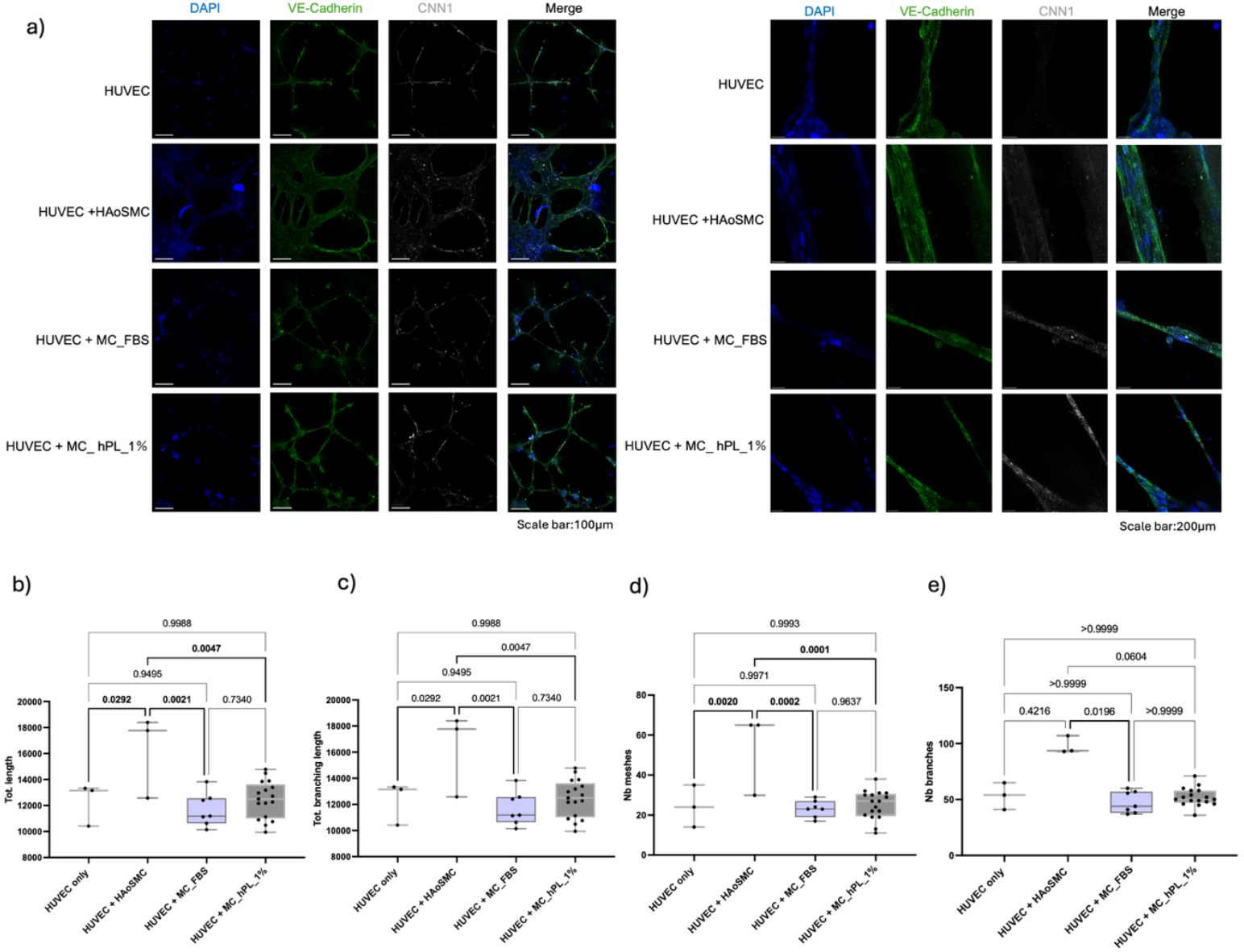
Vascular mural cell function assessment by endothelial angiogenic potential modulation effect. **a)** Representative immunofluorescent analysis of tube formation assay. Human umbilical venous endothelial cells (HUVEC) were co-cultured (5:1 ratio) with either day 13 human pluripotent stem cell (hiPSC) derived mural cells (MC) differentiated under human platelet lysate 1% (MC_hPL) or fetal bovine serum 10% (MC_FBS) supplemented conditions; or with human aortic smooth muscle cells (HAoSMC) as a positive control. HUVEC only cultures served as negative control. The left panel portrays the overall network formation (scale bar:100μm), while the right panel represents a magnified view of the same conditions (scale bar : 200μm) (n=3) . DAPI: 4′,6-diamidino-2-phenylindole. CNN1: Calponin 1. **b)-e)** Quantitative analysis of tube formation assay using Angiogenesis Analyzer in image J. Statistical analyses were performed using one-way ANOVA with Tukey’s post-hoc test and multiple comparison analysis (alpha value = 0,05) (n≥3). Significant differences (p<0,05) are indicated by **bold** parenthesis. **b)**Total Length (Tot. length) : Sum of all the branches and segments in the network. **c)** Total branching length (Tot. Branching length) cumulative length of all branches. **d)** Number of meshes (Nb meshes): indicator of network complexity by counting closed loops in the network. **e)** Number of Branches (Nb branches): indicator of sprouting potential.

Nevertheless, although structural intricacy seemed lower for both FBS and hPL 1% compared to the positive control, the number of branches in hPL 1% derived MC group were equivalent to HAoSMC; while significantly reduced in FBS derived MC group (p=0.0196) (Figure 5e). This suggests a slightly higher angiogenic branching potential in hPL 1% MC coculture compared to FBS condition. Across all angiogenic measurements, although no statically significant differences were observed between FBS and hPL1% derived MC, the overall data distribution indicated higher median values for hPL1% group compared with FBS (Figure 5b-e and Supplementary Figure S1).

## Discussion

Human platelet lysate has been widely used as a robust alternative to replace animal derived serum in biological studies using stem cells[5,23,24] and 3D organoid development [4–6,11]. Although previous studies have validated the use of hPL in the differentiation of diverse cell types, to our knowledge, this is first report evidencing hPL as an effective serum-free alternative for the differentiation of MC from hiPSC, yielding over 80% MC (Figure 2b) with hPL 1% supplementation, comparable to FBS differentiation efficiency. From the perspective of medical resource utilization, the development of new applications for hPL as shown in this study contributes to the realization of sustainable regenerative medicine [25].

Although there is still no consensus on specific MC markers [26,27], bulk RNA-sequencing profile comparison between FBS, hPL1% and hPL10% showed clear genetic differentiation between the groups (Figure 4a and 4d). Both FBS and hPL1% showed mural cell commitment with conserved co-expression of known pericyte marker PDGFR-β, NG2 and NGFR[21]. Interestingly, although other pericyte markers like DLK-1 and NT5E showed a decreased expression in hPL 1% compared to FBS, concomitant increased co-expression of contractile mural cell markers (TGLN, ACTA2 and specifically CNN1 and MYH11) may suggest a shift towards a more mature mural cell or partial smooth muscle phenotype in hPL1% derived MC compared to FBS[28,29]. This was further supported by gene ontology functional analysis showing enrichment in of pathways related to muscle contractility (Figure 4b) emphasizing cytoskeletal organization and mechanical integrity of muscle tissue in hPL1%, while in FBS enriched pathways were related to extracellular matrix (ECM) regulation and cell adhesion (Figure 4c) which is supported by previous reports on muscle pericyte phenotype[30]. These results also indicate that although both FBS and hPL1% lead to MC differentiation, there are differences in phenotype of the MC being differentiated. The encoding of smooth muscle contractile genetic expression in hPL1% is an important finding as it indicates a differential MC subphenotype expression closer to vSMC in hPL1% and pericytes for FBS derived MC [30,31].

On the other hand, hPL10% promoted a high expression of ACTA2, ACTN2, ACTC1, TNNI1, TNNT2, MYBPC3, MYH7, MYH11, MYL7 and CKM compared to hPL1% and FBS, compatible with a mixed cardiac and smooth muscle lineage[32] which explains the increased cardiomyocyte differentiation seen in this group. Of note, hPL1% evidenced a more discrete but concomitantly upregulated expression of some of these genes (Figure 4d and 4e), suggesting increasing concentration of hPL may enhance the differentiation of hiPSC away from mural phenotype towards cardiomyocyte, while lower concentrations of hPL might reinforce mural cell differentiation with a slightly residual cardiomyocyte differentiation pathway as evidenced in the transcriptional profile. However further studies using single RNA sequencing to avoid bias of residual cardiomyocyte differentiation expression in hPL 1% cell cultures are required, to further characterize the real gene expression differences between the MC yielded by FBS or hPL1% culture.

Regarding functional evaluation of the differentiated MC, it is well known that during angiogenesis endothelial cells actively recruit MC by secretion of the growth factor PDGFB to stabilize the growing vasculature[10]. Both pericytes and vSMC have diverse functions and therefore are identified with diverse protein markers. Pericytes line capillaries and venules, whereas vSMCs encircle arteries, arterioles, and veins[10]. In this context, MC differentiated with hPL1% proved to have comparable functionality to FBS in angiogenic processes settings (Figure 5a), with a slightly increase sprouting capacity comparable to mature vSMC control (HAoSMC) (Figure 5e). Nevertheless, both groups exhibited overall reduced functionality in most of the tube formation assay parameters compared HAoSMC (Figure 5b-e, Supplementary Figure S1), further suggesting that the differentiated MC are still in immature state[29].

Vascularization of hiPSC derived tissues and 3D organoids represents one of the biggest challenges in tissue engineering [33]. The development of clinical safe, vascularized organoids from hiPSC is a critical step towards translating regenerative medicine into clinical grade tissue engineering, with potential therapeutic applications in humans[34]. This study presents hPL 1% as a promising alternative to FBS for the differentiation of functional MC (PDGFR β +/CNN1+) from hiPSC. Vascularized organoids generated under xeno-free conditions could overcome one of the key bottlenecks in regenerative medicine; the ability to deliver functional, perfusable tissues that integrate seamlessly with host vasculature[35]. Such constructs hold promise not only for disease modeling and drug discovery, but also for future transplantation therapies, where the presence of stable mural cells is expected to enhance graft survival, maturation, and long-term function[36,37]. By enabling the production of clinically compatible vascular mural cells, our findings may accelerate the translation of hiPSC-derived vascularized organoids from the laboratory to therapeutic applications in patients.

Several limitations must be considered for future translational implementation. First, the biological variation of the hPL source, along with the lack of standardized protocols for processing, storage and handling represents a limitation of both this study and more importantly the overall hPL implementation potential[6]. This variability may directly impact the differentiation efficiency for all cell types, as observed in our protocol, especially with high hPL concentrations (Figure 2b-e). Storage, handling and freeze and thaw cycles and their effect on the hPL stability are variables that need further study. Furthermore, FBS is not the only animal derived product used this protocol. Therefore, to achieve a completely xeno-free MC differentiation system, further efforts to replace elements like Matrigel in hiPSC derived 3D organoid tissues for translational medicine must be considered.

## Conclusion

Overall, this study demosntrates that hPL serves as a robust alternative to FBS for the differentiation of MC from hiPSC, marking a technological breakthrough toward the establishment of a xeno-free, clinically compliant MC induction system. This advancement not only ensures safer and ethically sustainable MC derived hiPSC differentiation, but also accelerates the clinical translation of regenerative tissues, carrying significant clinical and social value as the development of clinical grade differentiation systems ultimately aims to enhance patient outcomes and broaden therapeutic and curative possibilities in regenerative medicine.

## Data availability statement

All data generated or analyzed during this study is available in accordance by request to the corresponding author. Bulk RNA sequencing data is available in the following link: https://drive.rdm.kyoto-u.ac.jp/index.php/s/pB9JeSEzcHmQBik

## Abbreviations

MC: Vascular mural cells
FBS: fetal bovine serum
hPSCs: human pluripotent stem cells
hPL: human platelet lysate
GMP: Good Manufacturing Practice
hiPSCs: human induced pluripotent stem cells
EDTA: ethylenediaminetetraacetic acid
PBS: phosphate-buffered saline
BMP4: bone morphogenetic protein 4
HUVEC: human venous umbilical endothelial cells
HAoSMC: human aortic smooth muscle cells
EGM-2: endothelial Cell Growth Medium 2
SMGM: smooth muscle cell growth medium 2
VEGF: vascular endothelial growth factor
PE: phycoerythrin
cTnT: anti–cardiac troponin
APC: allophycocyanin
DPBS: Dulbecco’s phosphate-buffered saline
vSMC: vascular smooth muscle cell
ACTA2: anti alpha smooth muscle actin
CNN1: calponin 1
GO: gene ontology
MYH11: myosin heavy chain 11
TAGLN: transgelin
ECM: extracellular matrix

## Acknowledgements

We sincerely thank Dr. Masaya Hagiwara for his generous support and guidance.

## Authors contributions

LY.GT. and H.M. conceptualized and designed the research studies. LY.GT. conducted experiments and acquired data. LY.GT., W.M. and H.M. analyzed the data. LY.GT. and H.M. wrote the manuscript. H.M. secured funding. K.M. and H.M. supervised this project. All authors reviewed and approved the final manuscript.

## Usage of generative AI and AI-assisted technologies in the manuscript writing process

During the preparation of this manuscript, the authors used OpenAI tools to improve readability during proof reading. Authors take full responsibility for the final version of the publication.

## Materials & Correspondence

Correspondence and material requests should be addressed to the corresponding author Hidetoshi Masumoto.

