## Supplementary Figure 1 for "Human platelet lysate drives clinically compliant generation of vascular mural cells from human pluripotent stem cells"

### Title

Hidetoshi Masumoto, MD, PhD

Department of Cardiovascular Surgery, Graduate School of Medicine, Kyoto University

54 Kawara-cho, Shogoin, Sakyo-ku, Kyoto, 606-8507, Japan

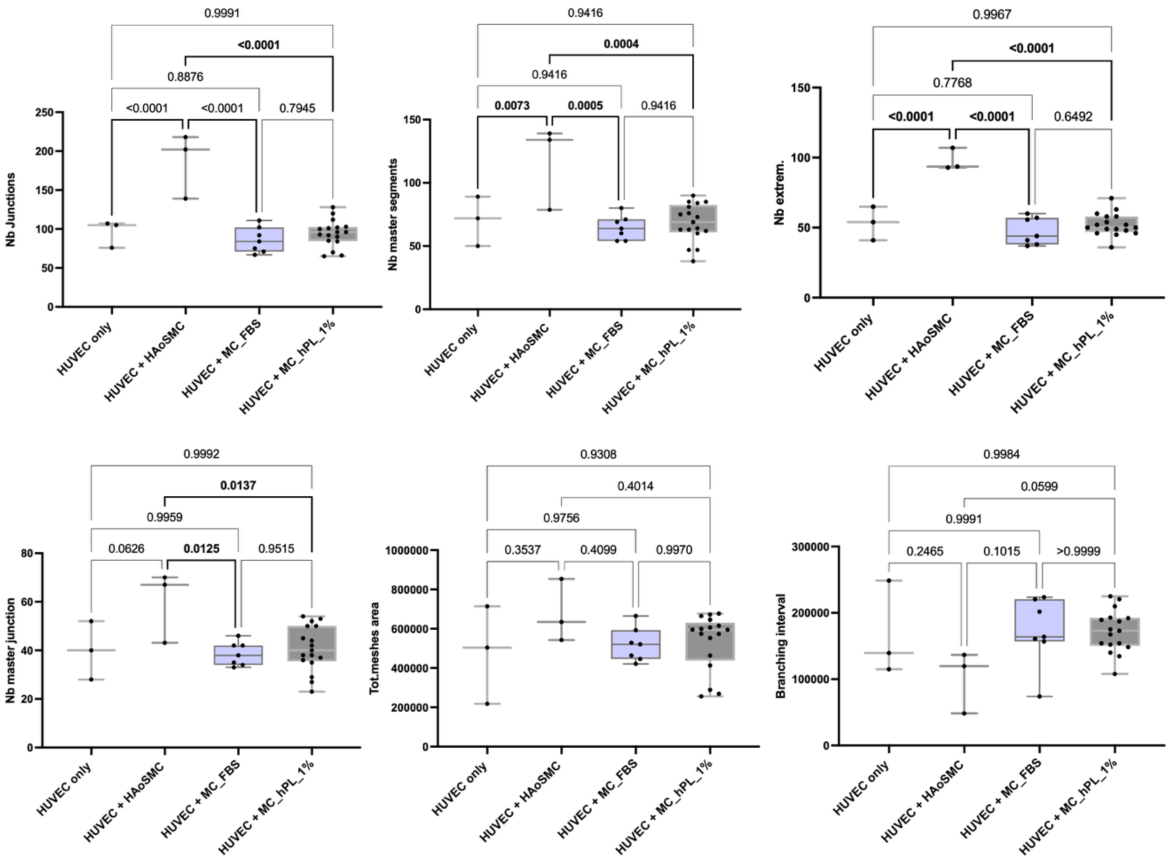


**Supplementary Figure 1. Additional evaluation parameters of vascular mural cell function through modulation of endothelial angiogenic potential**

Quantitative analysis of tube formation assay using Angiogenesis Analyzer in image J. Human umbilical venous endothelial cells (HUVEC) were co-cultured (5:1 ratio) with either day 13 human pluripotent stem cell (hiPSC) derived mural cells (MC) differentiated under human platelet lysate 1% (MC_hPL) or fetal bovine serum 10% (MC_FBS) supplemented conditions; or with human aortic smooth muscle cells (HAoSMC) as a positive control. HUVEC only cultures served as negative control. Statistical analyses were performed using one-way ANOVA with Tukey´s post-hoc test and multiple comparison analysis (alfa value = 0,05) (n≥3). Significant differences (p<0,05) are indicated by bold parenthesis. Nb Junctions: number of junctions (top left). Nb master segments: number of master segments (top middle). Nb extrem.: number of extremities (top right). Nb master junction: number of master junction (bottom left). Tot.meshes area: total of meshes area (bottom middle).
